# Tricellular junction proteins promote disentanglement of daughter and neighbour cells during epithelial cytokinesis

**DOI:** 10.1101/309922

**Authors:** Zhimin Wang, Floris Bosveld, Yohanns Bellaïche

## Abstract

In epithelial tissue, new cell-cell junctions are formed upon cytokinesis. To understand junction formation during cytokinesis, we explored in *Drosophila* epithelium, *de novo* formation of tricellular septate junctions (TCJs). We found that upon midbody formation, the membranes of the two daughter cells and of the neighbouring cells located below the adherens junction (AJ) remain entangled in a 4-cell structure apposed to the midbody. The septate junction protein Discs-Large and components of the TCJ, Gliotactin and Anakonda accumulate in this 4-cell structure. Subsequently, a basal movement of the midbody parallels the detachment of the neighbouring cell membranes from the midbody, the disengagement of the daughter cells from their neighbours and the reorganisation of TCJs between the two daughter cells and their neighbouring cells. While the movement of midbody is independent of the Alix and Shrub abscission regulators, the loss of Gliotactin or Anakonda function impedes both the resolution of the connection between the daughter-neighbour cells and midbody movement. TCJ proteins therefore control an additional step of cytokinesis necessary for the disentanglement of the daughter cells and their neighbours during cytokinesis.

## Introduction

Cytokinesis is the final step of cell division ensuring the physical separation of the two daughter cells after chromosome segregation. During cytokinesis, the constriction of the actomyosin contractile ring drives the ingression of the cleavage furrow until a narrow intercellular bridge remains, the midbody. The midbody serves as a platform for the recruitment of the abscission machinery, which eventually separates the two daughter cells via ESCRT-III-mediated membrane fission (for review, Glotzer, 2017 and Mierzwa and Gerlich, 2014). The midbody has also been shown to be involved in cell division orientation, cell fate specification, dorsal-ventral axis specification and ciliogenesis as well as tumorigenesis (Bernabé-Rubio et al., 2016; Dubreuil et al., 2007; Ettinger et al., 2011; Kuo et al., 2011; Singh and Pohl, 2014).

In invertebrate and vertebrate monolayered epithelial tissues, cytokinesis is tightly coordinated with the establishment of the apical-basal cell polarity at the newly formed interfaces between the daughter cells and their neighbouring cells (for review Herszterg et al., 2014; Higashi and Miller, 2017). This coordination is proposed to be critical for the maintenance of the epithelial tissue barrier function and involves an interplay between the dividing cell and its neighbouring cells located on each side of the cytokinetic ring (thereafter referred to as the neighbour cells). In all monolayered animal epithelial tissues examined so far, with the exception of the *Drosophila* embryo epithelium (Guillot and Lecuit, 2013), the cytokinetic ring deforms the membranes of both the dividing cell and the neighbouring cells at its rim leading to their co-ingression and the formation of a 4-cell structure apposed to the midbody (Firmino et al., 2016; Founounou et al., 2013; Herszterg et al., 2013; Higashi et al., 2016; Jinguji and Ishikawa, 1992; Lau et al., 2015; Morais-De-Sá and Sunkel, 2013; Pinheiro et al., 2017; Reinsch and Karsenti, 1994). Following midbody formation, the 4-cell structure can either resolve to form a “daughter-daughter” interface or a “neighbour-neighbour” interface.

The formation of a daughter-daughter interface requires actomyosin contractility in the neighbouring cells (Firmino et al., 2016; Herszterg et al., 2013). It is associated with a mechano-sensing mechanism triggered in the neighbouring cell upon cytokinetic ring constriction (Pinheiro et al., 2017). Cytokinetic ring constriction promotes a local membrane elongation resulting in a local decrease in E-Cadherin level at the rim of the cytokinetic ring, thereby producing actomyosin flows in the neighbouring cells to ensure the juxtaposition of the daughter cell adherens junctions (AJs). In turn, the formation of the midbody promotes an Arp2/3-dependent F-actin flow within the dividing cell leading to the withdrawal of the neighbouring cell membranes at the apical AJs of the cells to establish a daughter-daughter cell AJ (Herszterg et al., 2013). The formation of a neighbour-neighbour interface in epithelial cells of the gastrulating *Xenopus* embryo was shown to depend on a distinct mechanism, which is mediated by Vinculin and that reinforces AJ stabilization at the rim of the cytokinetic ring (Higashi et al., 2016). As shown in both invertebrate and vertebrate epithelial tissues, the formation of a daughter-daughter or neighbour-neighbour interface, as well as the apical-basal position of the *de novo* AJ, contribute to the final arrangement of the cells within the tissue and the overall organization and dynamics of the tissue during development (Firmino et al., 2016; Herszterg et al., 2013; Lau et al., 2015; Morais-De-Sá and Sunkel, 2013).

The co-ingression of both the dividing cell and neighbouring cell membranes leading to the creation of a 4-cell structure upon midbody formation raises the question of the mechanisms by which the dividing cell ensures its disentanglement from its neighbouring cells while maintaining epithelial barrier function. The *de novo* AJ and tight tricellular junction formation upon cytokinesis and the resolution of the 4-cell structure leading to the formation of a neighbour-neighbour interface have recently been described in *Xenopus* epithelial tissues (Higashi et al., 2016). Yet, the mechanisms are not fully understood when a daughter-daughter interface is formed.

To explore the mechanism by which a 4-cell structure formed around the midbody resolves to lead to the formation of a daughter-daughter interface, we undertook a detailed analysis of the *de novo* bicellular and tricellular septate junction (TCJ) formation in dividing cells of the *Drosophila* dorsal thorax epithelium tissue, where all divisions result in the formation of a daughter-daughter cell interface. Doing so, we uncovered that the daughter and neighbouring cells remain tightly entangled long after midbody formation. The disentanglement of the 4-cell structure occurs as the midbody moves to a more basal position; this midbody movement being associated with the *de novo* TCJ formation between the two daughter cells and each of their neighbouring cells. Furthermore, we uncovered that the neighbour cell disengagement from the midbody and the midbody movement are impaired in the absence of the Gliotactin (Gli) and Anakonda (Aka) TCJ proteins.

## Results

### The septate junction marker Dlg labels a 4-cell structure upon bicellular AJ and SJ formation

During mitosis, the *de novo* AJ formation between the two daughter cells is tightly coupled with the ingression of the cytokinetic ring and the formation of the midbody (Fig. 1A; Firmino et al., 2016; Founounou et al., 2013; Herszterg et al., 2013; Higashi et al., 2016; Jinguji and Ishikawa, 1992; Lau et al., 2015; Morais-De-Sá and Sunkel, 2013; Pinheiro et al., 2017; Reinsch and Karsenti, 1994). To better understand the mechanisms of junction formation in *Drosophila* epithelia, we studied the dynamics of *de novo* septate junction (SJ) formation during cytokinesis and compared it to the formation of the AJ in the epithelial cells of the pupal dorsal thorax.

**Figure 1.**
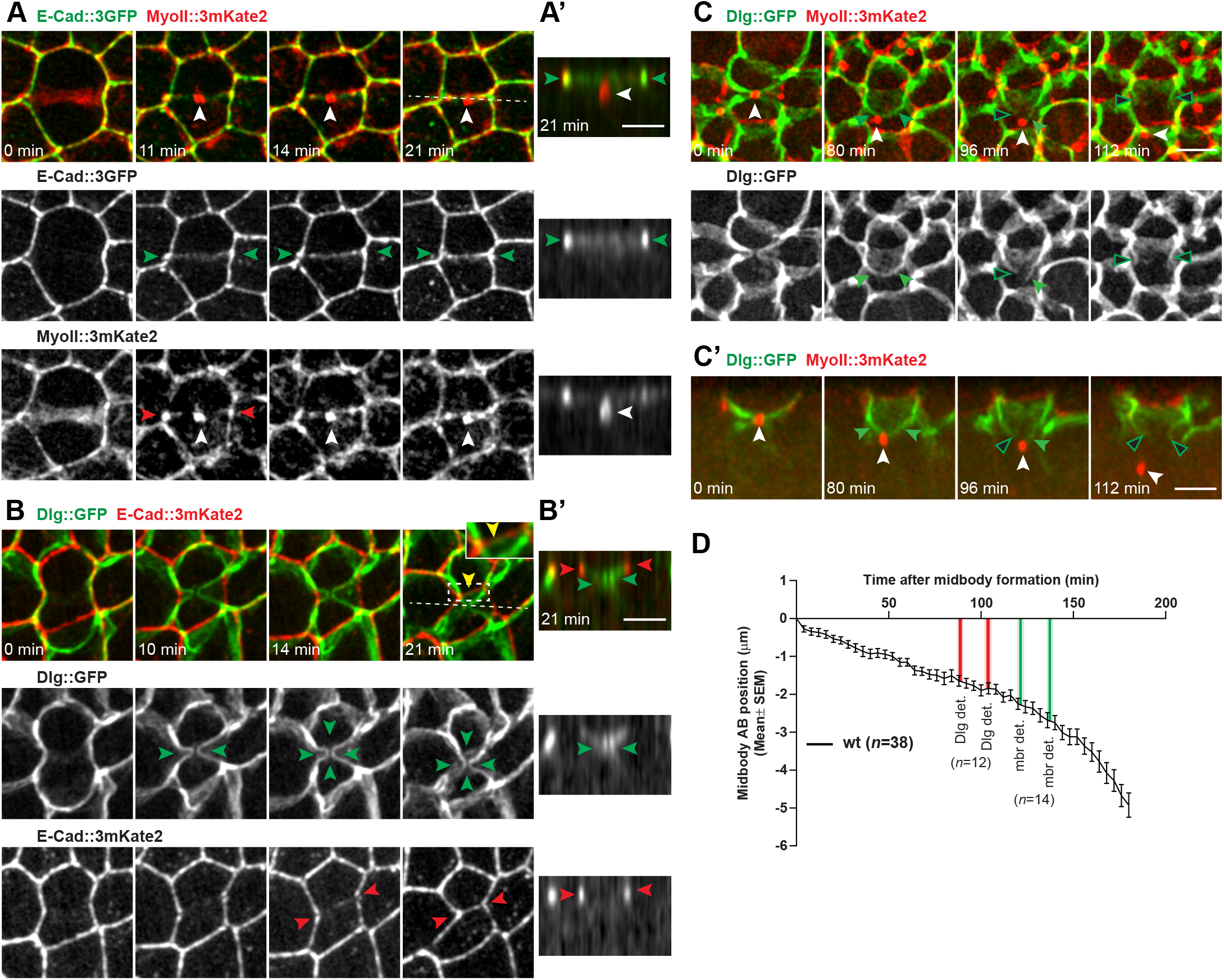
Cytokinetic ring constriction induces the formation of a 4-cell structure. (A-A’) Confocal z-section projections of E-Cad::3GFP (green in the top panel, white in the middle panel) and MyoII::3mKate2 (red in the top panel, white in the bottom panel) distributions in the dividing cell and its neighbours during cytokinesis (A). Apical-basal section of the cell in A at t=21 min along the dashed line (A’). Red arrowheads, MyoII::3mKate2 accumulation in the neighbouring cells from mid-constriction onwards; green arrowheads, newly forming AJ; white arrowheads, midbody. (B-B’) Confocal z-section projections of Dlg::GFP (green in the top panel, white in the middle panel) and E-Cad::3mKate2 (red in the top panel, white in the bottom panel) distributions in the dividing cell and its neighbours during cytokinesis (B). Yellow arrowhead, position of the bicellular SJ formed just below the AJ and between the two daughter cells (dashed box magnified in the top right); green arrowheads, 4-cell structure formed more basally at the interface between the two daughters and two neighbour cells; red arrowheads, forming AJ between the two daughter cells. Apical-basal section of the cell in B at t=21 min along the dashed line (B’). (C-C’) Confocal z-section projections (C) and apical-basal sections (C’) of Dlg::GFP (green in the top and bottom panels, white in the middle panel) and MyoII::3mKate2 (red in the top and bottom panels) distributions in the dividing cell and its neighbours during cytokinesis. White arrowheads, midbody; green arrowheads, Dlg::GFP positive strands forming between the apical SJ and the midbody; open green arrowheads, Dlg::GFP positive strands not apposed to the midbody. (D) Graph of the apical-basal position of the midbody (Mean±SEM) relative to its initial apical-basal position in wt cells. The red versus green vertical bars indicate the timing at which the first and second Dlg::GFP strands (91±3 and 106±4) versus PH::GFP labelled membrane strands (mbr, 120±4 and 140±3.5) detach from the midbody (det.: detachment, i.e. the Dlg::GFP or PH::GFP signal is not closely apposed to the midbody). The shaded domain corresponds to the SEM of the timing of each detachment event. Scale bars: 5 μm. *n*, cell numbers (D).

To this end, we imaged the dynamics of a functional Dlg::GFP, as a marker of the SJ, and of E-Cad::3mKate2 as a marker of the AJ (Fig. 1B; Movie 1). In parallel, we analysed the dynamics of Dlg::GFP and MyoII::3mKate2 (Sqh), which marks the cytokinetic ring and the midbody (Fig. 1C and Fig. S1; Movie 2). The SJ and AJ of the dividing cell and its neighbours deform concomitantly with early ring constriction (Fig. 1B, t=0 min). Around mid-constriction, MyoII accumulation in the neighbouring cells promotes the juxtaposition of the ingressing membrane at the level of the AJ as previously reported (Fig. 1A, Herszterg et al., 2013; Pinheiro et al., 2017), while the Dlg::GFP labelled SJ remains far apart, forming an ingression connecting the two daughter cells and their two neighbouring cells on each side of ring (Fig. 1A and B, at t=11 and 10 min, respectively, green arrowheads). As the daughter-daughter cell AJ is established upon midbody formation (Herszterg et al., 2013), Dlg::GFP begins to accumulate just below the AJ decorating the newly forming bicellular SJ (Fig. 1B, yellow arrowhead at t=21 min and inset). In addition, a pool of Dlg::GFP is localized more basally (Fig. 1B’,C’). More precisely, Dlg::GFP remains present at the interfaces of two daughter cells and their two neighbours; these interfaces being tightly apposed to the MyoII positive midbody that is located below the bicellular AJ and SJ formed between the two daughter cells (Fig. 1A’, C’, green and red arrowheads, respectively).

Together these data confirmed that the *de novo* formation of a bicellular AJ is associated with midbody formation. Furthermore, these findings show that (i) a bicellular Dlg::GFP SJ localized just below the AJ is assembled between the two daughter cells concomitantly to AJ formation and (ii) Dlg::GFP is also enriched at the interfaces of the four cells involved in the cytokinesis process: the two daughters and the two neighbouring cells located on either side of the cytokinetic ring; hereafter referred to as the “4-cell structure” (Fig. 1B, C).

### The resolution of the 4-cell structure is associated with a midbody basal movement

To analyze the fate of the 4-cell structure, we prolonged our time-lapse analyses of the MyoII::3mKate2 and Dlg::GFP dynamics (Fig. 1C, D; Movie 2). Since the 4-cell structure forms after midbody formation, we timed from thereon all events relative to midbody formation and imaged these markers in cells that are slightly tilted relative to the apical-basal axis to facilitate the exploration of their 3-dimensional (3D) organization and dynamics in 2-dimentional (2D) projections. These analyses were also performed for other SJ markers Lachesin::YFP, α-ATPase::GFP and Neuroglian::GFP, the dynamics of which mirror the ones described for Dlg::GFP (Fig. S2A and not shown).

Following the distribution of Dlg::GFP and MyoII::3mKate2 for more than 3 hours upon midbody formation revealed that the midbody exhibits a stereotypical movement to a more basal position and away from its site of formation (Fig. 1C-D, Fig. S2B). During this period, Dlg::GFP is modestly enriched along the apical-basal bicellular interface formed between the two daughter cells; this daughter-daughter interface adopting a triangular shape (Fig. 1C, t=80 min). More interestingly, as the midbody descends the Dlg::GFP signal remains highly enriched along the position between the two daughter cells and each neighbouring cell, forming two Dlg::GFP positive strands. The apical tip of each Dlg:GFP positive strand is located at the AJ vertex formed by the 2 daughter cells and their neighbouring cells, while the basal tip of each Dlg::GFP strand is tightly apposed to the midbody. As the midbody is displaced more basally, the Dlg::GFP positive strands elongate until the basal tip of one of the two Dlg::GFP positive strands loses its tight apposition to the midbody (Fig. 1C,C’, t=96 min and on average 91±3 standard error of the mean, SEM, *n*=12, Fig. 1D, Fig. S2B). This event is then followed by the loss of the tight apposition of the second Dlg::GFP positive strand to the midbody (Fig. 1C,C’, t=112 min and on average 106±4 min, *n*=12, Fig. 1D, Fig. S2B). While the existence of a direct physical connection between the midbody and the tip of the Dlg::GFP positive strand remains to be established, we thereafter refer to this step as the “Dlg basal tip detachment”. Following the second strand detachment, the midbody continues to descend and moves away from its initial position (Fig. 1D, Fig. S2B) and each Dlg::GFP strand formed between the two daughter cells and one neighbouring cell adopts an apical-basal orientation (Fig. 1C, Movie 2). These analyses led us to hypothesize that (i) the neighbouring cells remain tightly connected to the midbody for an extensive period of time and that (ii) each of the Dlg::GFP positive strands, located between the two daughter cells and one neighbouring cell, is the prospective TCJ forming between the two daughter cells and each of their neighbouring cells.

### The daughter and neighbour cells are entangled along the membrane of the future tricellular junction

To test whether the midbody remains associated with both the daughter and the neighbouring cells, we specifically labelled either the daughter or the neighbouring cells in cells co-expressing MyoII::3mKate2 to follow the position of the midbody. The labelling of the two daughter cell cytoplasm using the F-Actin marker LifeAct::GFP confirmed that the MyoII::3mKate2 labelled midbody is localized between the daughter cells as it adopts a basal position and moves away from its site of formation (Fig. 2A). To better study whether the two neighbouring cells remain connected to the midbody, we imaged dividing cells surrounded by two neighbouring cells expressing the membrane marker PH::GFP in a MyoII::3mKate2 expressing tissue (Fig. 2B). This revealed that the membranes of the two neighbour cells remain tightly apposed to the midbody for more than 2 hours upon midbody formation (Fig. 2B). The PH::GFP and the MyoII::3mKate2 signals were then used to manually segment the membranes of the two neighbouring cells and the position of the midbody, respectively, to reveal their 3D organization. As suggested by the analysis of Dlg::GFP, the 3D reconstitution illustrated that the membranes of the two neighbouring cells ingress between the two daughter cells and remain apposed to the midbody (Fig. 2B’). Such organization perdures as the midbody adopts a more basal position. Strikingly, on average 2 hours after midbody formation (120±4 min, *n*=14), one of the two neighbouring cell membranes loses its tight apposition to the midbody. This first detachment is then followed by the detachment of the second neighbouring cell membrane (140±3.5 min, *n*=14) (Fig. 1E; Fig. 2B,B’; Fig. S2B). As the midbody is localized between the two daughter cells (Fig. 2A,B), this result supports the existence of a 4-cell structure connecting the two daughter and two neighbouring cells to the midbody that remains upon the formation of the apical daughter-daughter AJ interface.

The timing of neighbour cell membrane detachment from the midbody indicates that the detachment of the Dlg::GFP basal strand tips occurs prior to membrane detachment. Accordingly, time-lapse analyses of PH::GFP expressed in one of the two neighbouring cells in a Dlg::TagRFP and MyoII::mKate2 tissue confirmed that the membrane of the neighbouring cells remained associated with the midbody as the midbody became localized basally and away from the basal tip of the Dlg::TagRFP labelled strands. This shows that the neighbouring cell membranes remain apposed to the midbody, even after the Dlg::GFP basal tips detached from the midbody (Fig. 2C,C’).

**Figure 2.**
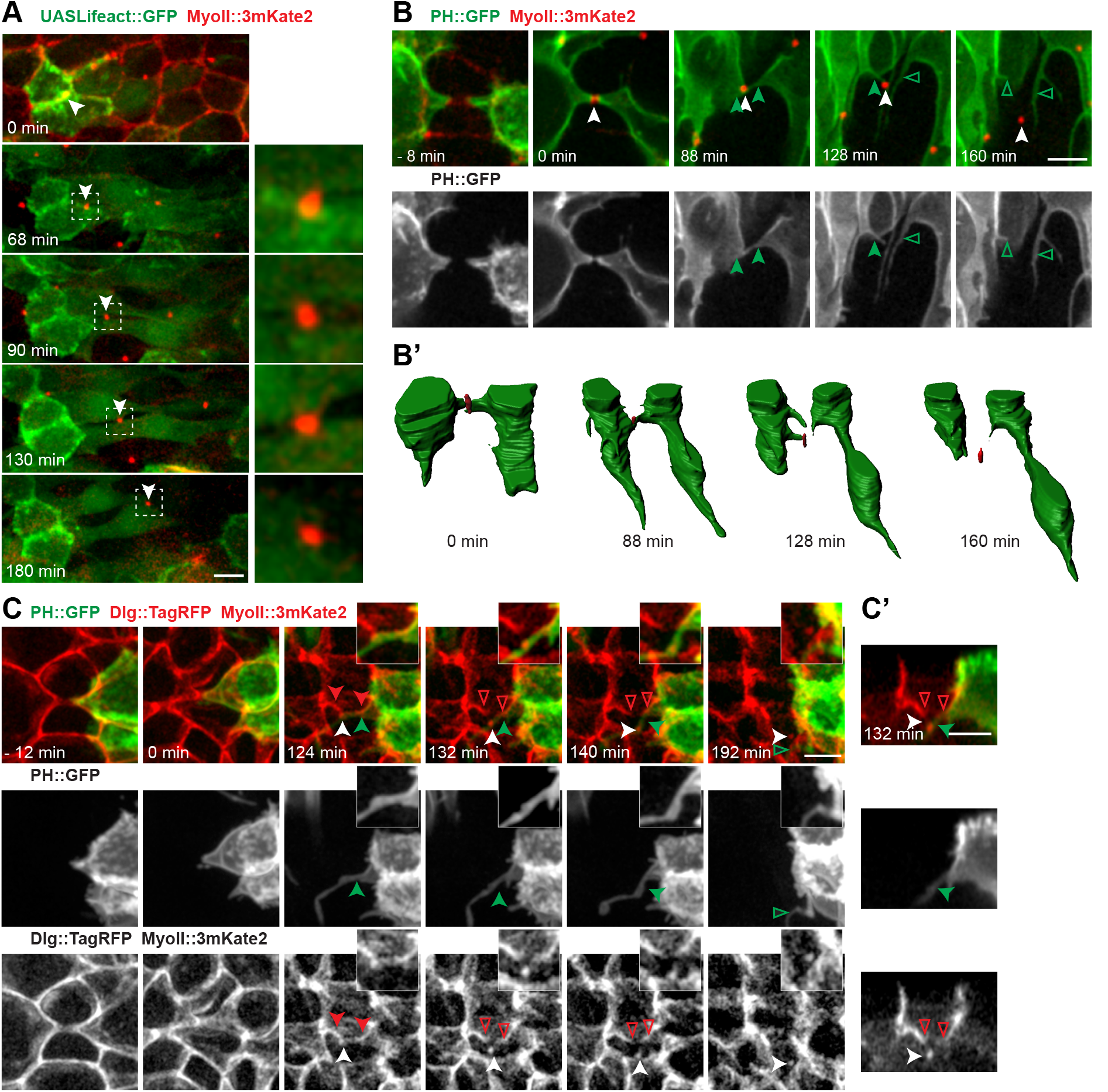
The 4-cell structure disentangles during midbody descend. (A) MyoII::3mKate2 (red) distribution and LifeAct::GFP (green) distribution in the 2 daughter cells during cytokinesis. A full apical-basal projection of the confocal z-section is shown for the LifeAct::GFP signal, whereas, to better highlight the MyoII labelled midbody and to avoid the apical MyoII cortical signal, a 1 μm confocal z-section projection around the MyoII::3mKate2 marked midbody is shown. White arrowheads, midbody. Dashed box regions, insets magnified in the right panels. (B-B’) PH::GFP (green in the top panel, white in the bottom panel) and MyoII::3mKate2 (red in the top panel) distributions in the 2 neighbours of the dividing cell. A 1 μm z-section projection around the midbody is shown. White arrowhead, midbody; green arrowheads, membranes of the neighbouring cells extending to the midbody; open green arrowheads membranes detached from the midbody. The lower panels show the 3D reconstructions of the neighbouring cell contours (green) and of the midbody of the dividing cell (red) (B’). (C-C’) PH::GFP (green in the top panel, white in the middle panel), MyoII::3mKate2 and Dlg::TagRFP (red in the top panel, white in the bottom panel) distributions during cytokinesis. PH::GFP is specifically expressed in one of the two neighbouring cells. A full projection of the confocal z-section is shown. Apical-basal section of the cell in C at t=132 min (C’). White arrowheads, midbody; green and red arrowheads, membrane of the neighbouring cell and Dlg::TagRFP positive strands extending to the midbody, respectively; open green and open red arrowheads, membrane of the neighbouring cell and Dlg::TagRFP positive strands not apposed to the midbody, respectively. Scale bars: 5 μm.

To explore whether TCJs form along the Dlg::GFP strands connected to the midbody, we recorded the dynamics of Dlg::TagRFP, MyoII::3mKate2 and the TCJ protein Gli;::YFP (Fig. 3A,B; Movie 3). Dlg::TagRFP and Gli::YEP time-lapse movies revealed that Gli:YFP signal is enriched at the Dlg positive strands (Fig. 3A,A’). Accordingly, the tracking of the MyoII::3mKate2 labelled midbody revealed that (i) Gli::YFP is enriched along two strands formed between the apical vertex of the two daughter cells and each neighbouring cell; (ii) each strand extends as the midbody moves more basally and eventually detaches from the midbody as observed for Dlg::TagRFP (Fig. 3B,B’; Movie 3). The dynamics of the TCJ were further confirmed by studying the dynamics of Aka::GFP, which mirors the ones of Gli::YFP (Fig. S3A,B).

**Figure 3.**
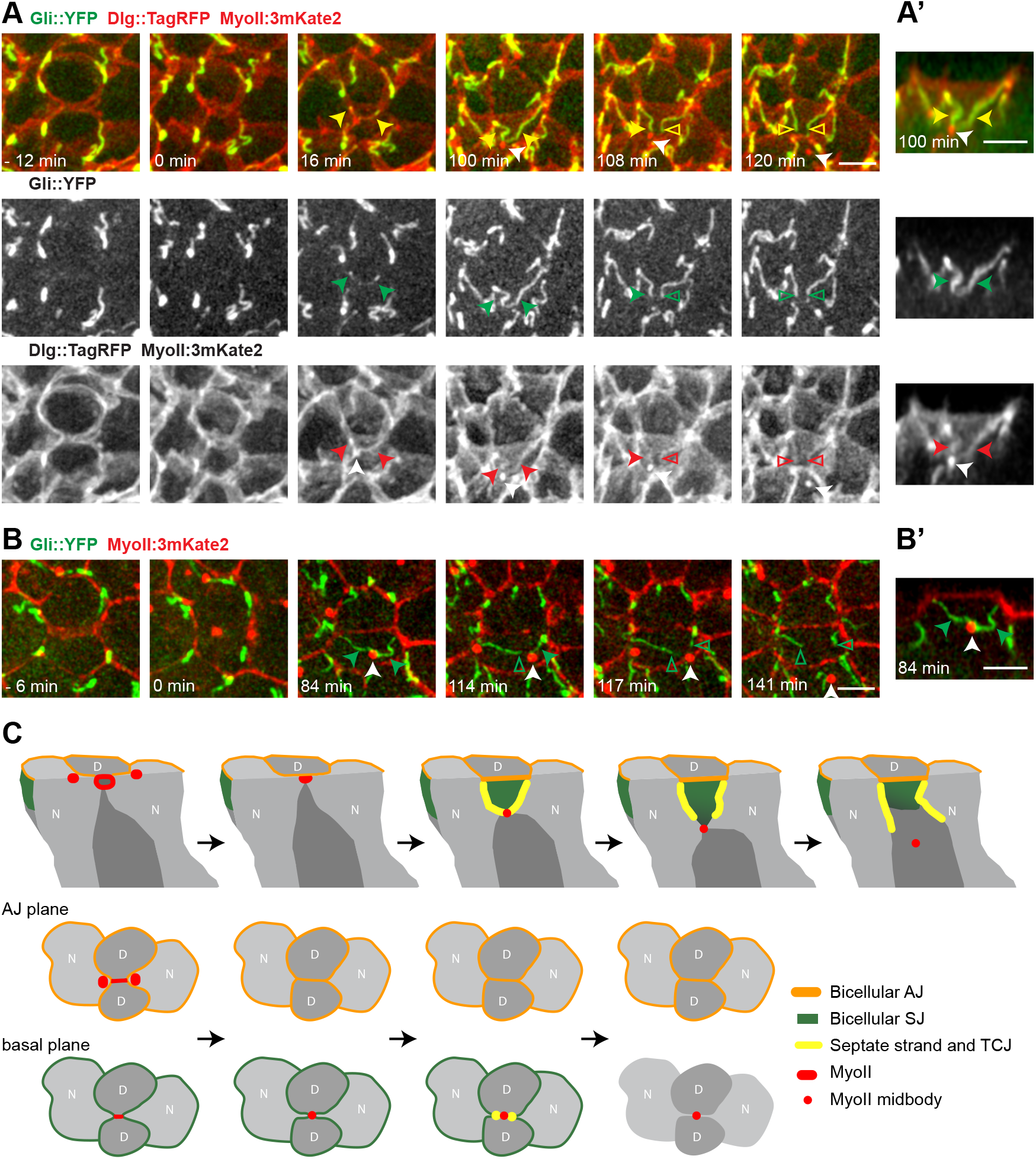
Dynamics of TCJ during cytokinesis. (A-A’) Gli::YFP (green in the top panel, white in the middle panel), Dlg::TagRFP and MyoII::3mKate2 (red in the top panel, white in the bottom panel) distributions during cytokinesis. A full projection of the confocal z-section is shown. Apical-basal section of the cell in A at t=100 min (A’). White arrowheads, midbody; green and red arrowheads indicate Gli::YFP and Dlg::TagFP positive strands, respectively, extending towards and apposed to the midbody; open green and red arrowheads, Gli::YFP and Dlg::TagFP positive strands, respectively, not apposed to the midbody; yellow arrowheads and open yellow arrowheads, Gli::YFP and Dlg::TagFP positive strands. (B-B’) Gli::YFP (green) and MyoII::3mKate2 (red) distributions during cytokinesis. A full projection of the confocal z-section is shown. Apical-basal section of the cell in B at t=84 min (B’). White arrowheads, midbody; green arrowheads, Gli::YFP strands extending towards and apposed to the midbody; open green arrowheads, Gli::YFP positive strands not apposed to the midbody. (C) Schematic of the dynamics of AJ (orang), SJ (green) and TCJ (yellow) formed between the 2 daughter (D, dark grey) cells and neighbour cells (N, light grey) in relation with midbody (red dot) movements during cytokinesis. Top panels: schematic cross sections showing apical-basal views along the bicellular junctions that is formed between the 2 daughters. Middle and bottom panels: schematic showing cross sections in the plane of the tissue at the levels of the AJ (middle panels) and the midbody (bottom panels). Upon cytokinetic ring contraction MyoII accumulates in the neighboring cells (red) and the midbody (red dot) is apically formed. Upon midbody initial basal movement, an apical daughter-daughter cell bicellular AJ and a bicellular SJ are formed between the two daughter cells, while more basally the daughter cells and the two neighboring cells form a 4-cell structure connected to the midbody. As the midbody moves more basally, a triangular interface is formed between the 2 daughter cells where the bicellular SJ components can be found. Septate and TCJ protein accumulate in the 2 strands formed between the apical vertex and the midbody. As the midbody is displaced more basally, the septate junction strands and TCJ strands detached from the midbody. Lastly, the membranes of the neighboring cells detached from the midbody and the TCJs adopt an apical-basal orientation. Scale bars: 5 μm.

Collectively our data highlight that (i) from an epithelial polarity point of view, (a) the apical daughter-daughter cell bicellular SJ is formed rapidly following midbody formation and (b) the two presumptive TCJs are formed during the midbody basal movement; their formation takes place within the 4-cell structure that later detaches from the midbody as the midbody moves further away from its initial position; (ii) from an epithelial organization point of view a 4-cell membrane structure apposed to the midbody is formed by the two daughter cells and the neighbouring cell membranes. This structure is tightly connected to the midbody and leads to an entanglement of the daughter cells with its neighbours. The resolution of this 4-cell structure starts by the detachment of one of the neighbouring cell membrane, on average 120 min after midbody formation; the second detachment taking place on average 20 min later (Fig. 1C,D; Fig. 2B; Fig. 3C; Fig. S2B).

### Characterization of the midbody movement and TCJ formation relative to cytokinesis steps

To further understand the mechanisms of TCJ formation during cytokinesis, we studied the dynamics of the basal midbody movement and of the formation of the TCJ relative to the dynamics of known markers or regulators of cytokinesis and abscission.

In agreement with previous work (Herszterg et al., 2013; Morais-De-Sá and Sunkel, 2013), we observed that an F-actin accumulation occurs around the midbody and that F-actin rapidly disappears after midbody formation (data not shown and see Herszterg et al., 2013). The F-actin accumulation preceeds the formation of the central spindle remnant marked by the Jupiter::GFP microtubule associated protein; the signal of which remains visible around the midbody for 25±1 min following midbody formation (*n*=14; Fig. 4A). The F-Actin accumulation around the midbody and the central spindle remnant are therefore unlikely to directly participate in the detachment of the forming TCJ from the midbody.

**Figure 4.**
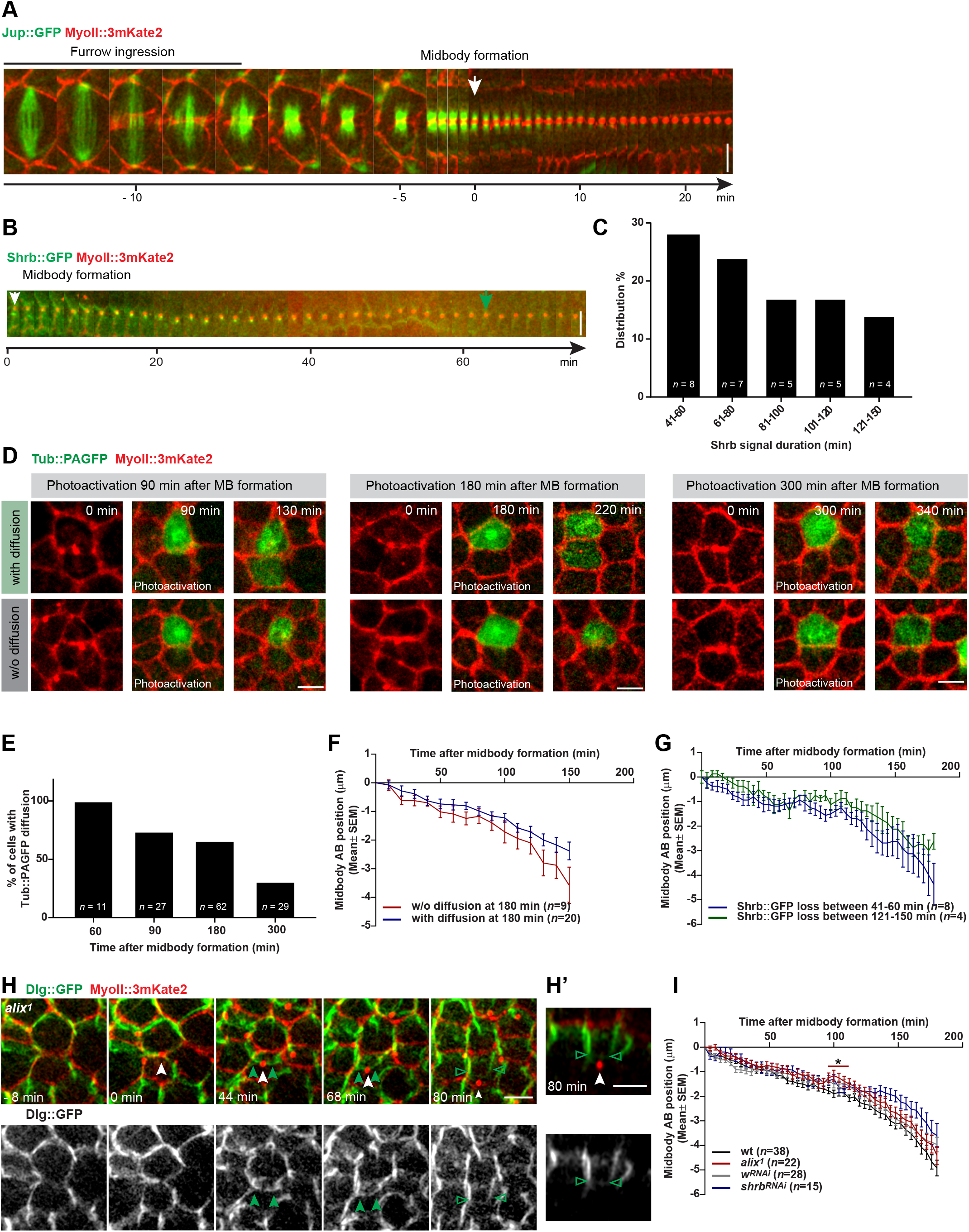
Daughter-neighbour disentanglement and midbody descent occur independently of abscission. (A-B) Kymographs of Jup::GFP (green) (A) and Shrb::GFP (green) (B) and MyoII::3mKate2 (red) (A, B) distributions during cytokinesis. White arrows, timing of midbody formation; green arrow in B, time at which Shrb::GFP is fully lost from the midbody (note that a weak Shrb::GFP signal is present on the daughter-daughter interface). (C) Histogram of the distribution of the time at which the Shrb::GFP signal disappears from the midbody. t=0 corresponds to midbody formation. (D) Tub::PAGFP (green) and MyoII::3mKate2 (red) time-lapse images at midbody formation (t=0), at the time of photoactivation of the Tub::PAGFP (at t=90 min left panels, at t=180 min middle panels and at t=300 min right panels) and 40 min post-photoactivation (t=130 min, t=220 min and t=340 min). In the upper panels, photoactivated Tub::PAGFP diffuses from the photoactivated daughter to the non-photoactivated daughter. In the lower panels, no diffusion between daughter cells is observed upon Tub::PAGFP photoactivation in one daughter cell. (E) Histogram of the fraction of cell divisions for which Tub::PAGFP diffuses between the two daughter cells upon Tub::PAGFP photoactivation of one daughter cell at different time points following midbody formation. (F) Graph of the midbody apical-basal positions (Mean±SEM) for which Tub::PAGFP diffuses between daughter cells (blue) and for cells for which Tub::PAGFP does not diffuse between daughters (red) upon photoactivation of Tub::PAGFP in one daughter cell 180 min after midbody formation. (G) Graph of the midbody apical-basal positions (Mean±SEM) for cells for which Shrb::GFP signal disappears between either 41-60 min (blue) or 121-150 min (green) after midbody formation. (H) Dlg::GFP (green in the top panel, white in the bottom panel) and MyoII::3mKate2 (red in the top panel) distributions in the dividing cell and its neighbours during cytokinesis in an *aliX^1^* clone (marked by the absence of nls::GFP, not shown). A full projection of the confocal z-section is shown. Apical-basal section of the cell shown in H at t=80 min (H’). White arrowheads, midbody; green arrowheads, Dlg::GFP strands extending and apposed to the midbody; open green arrowheads, Dlg::GFP strands not apposed to the midbody. (I) Graph of the midbody apical-basal position (Mean±SEM) in indicated genotypes. Clones were marked by the absence of nls::GFP for *aliX^1^* and the absence of cytoplasmic GFP for *W*^RN*Ai*^ and *shrb*^RN*Ai*^. Colored bar with asterisk indicates time-points for which the midbody positions are significantly different from control cells (*t*-test). Scale bars: 5 μm. *n*, cell numbers (C, E, F, G, I).

We then analysed whether the formation of the TCJ and the disengagement of the neighbouring cells is associated with abscission. We first analyzed the dynamics of the *Drosophila* CHMP4 orthologue ESCRT-III protein Shrub (Shrb) required for abscission (Capalbo et al., 2012; Eikenes et al., 2015; Glotzer, 2017; Matias et al., 2015). As expected, we found that Shrb::GFP localized at the midbody soon after its formation. The timing of Shrb::GFP disappearance from the midbody is on average 85±5 min following midbody formation (*n*=29, Fig. 4B). However, the timing of disappearance was variable from cell to cell and it ranges from 40 to 150 mins upon midbody formation (Fig. 4C). Shrb::GFP also remains weakly localized at the membrane interface between the two daughter cells upon disappearance of Shrb::GFP at the midbody (Fig. 4B). Importantly, we found that the dynamics of midbody movement are similar in cells for which Shrb::GFP signal disappears early or late after midbody formation (Fig. 4G).

To analyse whether the disappearance of the midbody Shrb::GFP signal might be related to abscission, we estimated the timing of abscission in this epithelial tissue. Abscission corresponds to the step at which the connection between the two daughter cells is severed. A lower bond for the abscission timing can therefore be determined by analysing the diffusion of small GFP tagged molecules between the two daughter cells. To determine the presence or absence of diffusion between the two daughter cells, we implemented a multiphoton photoactivation approach to label Tub::PAGFP in only one of the two daughter cells; multiphoton photoactivation being essential to avoid Tub::PAGFP activation in surrounding cells. By performing the analysis at different time-points following midbody formation, we found that photoactivated Tub::PAGFP always diffuses between daughter cells 60 min after midbody formation (*n*=11) (Fig. 4D,E). At 180 minutes after midbody formation, Tub::PAGFP still diffuses from one daughter to the other in the majority of the cells (66%, *n*=62). It is only 300 min (5 hours) after midbody formation that photoactivated Tub::PAGFP diffuses between daughter cells in the minority of cells (31%, *n*=29). These data indicate that 180 min after midbody formation, the majority of cells have yet to complete abscission; a result in agreement with our observation that the midbody is tightly associated with the two daughter cells when it is located basally (Fig. 2A). Since the neighbouring membrane detachment from the midbody occurs on average 140 min after midbody formation, this indicates that the disentanglement of the dividing cell and its neighbours can occur prior to abscission in the majority of the dividing cells. Accordingly, the basal descent of the midbody is not different in cells that undergo early or late abscission based on the Tub::PAGFP assay (Fig. 4F) or in cells where the Shrb::GFP signal disappears from the midbody at early or late time-points after midbody formation (Fig. 4G).

To further explore the relationship between abscission and the 4-cell disentanglement, we analysed whether the midbody movement depends on the activities of Shrb or Alix (ALG-2-interacting protein X), which are necessary for abscission (for review, Glotzer, 2017). Using both RNA interference (RNAi) approaches and loss of function alleles, we found that the midbody basal movements occur mainly as observed in controlled cells (Fig. 4H,I; Fig. S4). We therefore conclude that TCJ formation generally precedes abscision and that neither Shrb nor Alix are critical regulators of the basal mibody movement.

### Gli and Aka control daughter-neighbour disentanglement and midbody movement

So far, our results suggest that the basal midbody movement is associated with TCJ formation and the disengagement of the 4-cell structure. We therefore aimed to identify regulators of the midbody movement as an entry point to better understand the mechanisms of TCJ formation and neighbour cell disentanglement.

We first tested the role of AJ formation in midbody descent using α-Catenin RNAi. While the initial basal midbody descend is perturbed in *a-Cal*^RN*Ai*^ dividing cells, the midbody still moves basally and its final position is not significantly different compared to the ones of control dividing cells (Fig. 5A). As we observed that TCJ components are enriched in the two membranes connected to the midbody, we explored the roles of the TCJ proteins, Gli and Aka during cytokinesis. To this end, we imaged Dlg::GFP and MyoII::3mKate2 in either Gli or Aka RNAi knock-down conditions as well as in Gli and Aka loss of function alleles. Strikingly, whereas both Dlg:GFP positive strands have detached from the midbody in all *W*^RN*Ai*^ cells (*n*=16) 3 hours following midbody formation, they failed to detach from the midbody in 76% of cases (*n*=21) in *Gli*^RN*Ai*^ dividing cells and in 90% of cases in *aka*^RN*Ai*^ dividing cells (*n*=20) (Fig. 5B,D; Movie 4). Accordingly, we found that the midbody fails to move basally in *Gli*^RN*Ai*^ or *aka*^RN*Ai*^ divding cells; the common and most pronounced defects been observed 132 min after midbody formation (Fig. 5C,E). We confirmed these results using Gli and Aka loss of function alleles (Fig. S5). We could further show that Aka and Gli are required both in the dividing cell and the neighbouring cells for midbody movement (Fig. 5C, E). Together, we conclude that TCJs form as the midbody adopts a more basal position and that the TCJ proteins are critical for midbody movement and the disengagement of the neighbouring cells from the two daughter cells.

**Figure 5.**
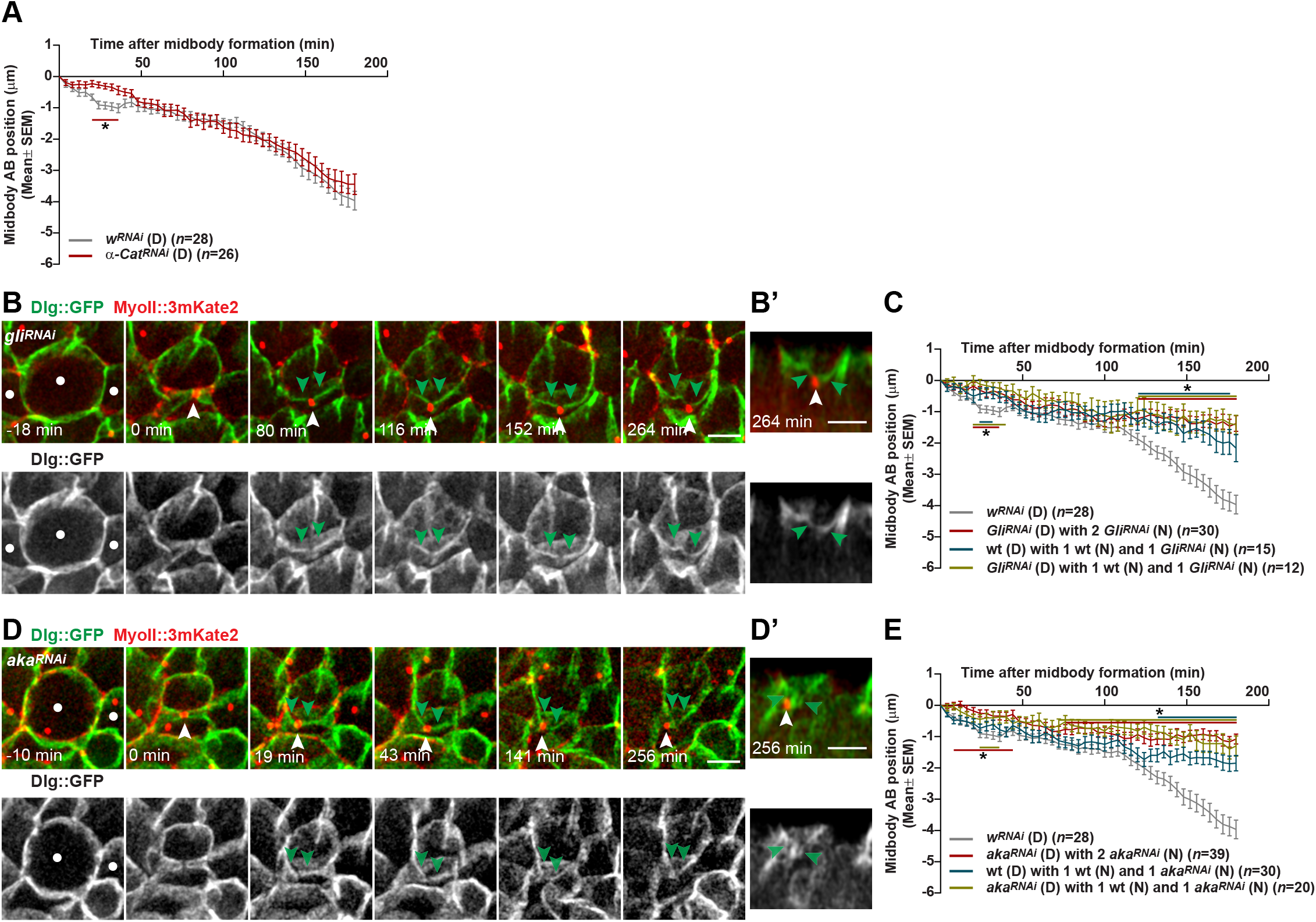
Midbody basal movement and daughter-neighbour disentanglement are abrogated when *de novo* TCJ formation is blocked. (A) Graph of the midbody apical-basal positions (Mean±SEM) in α-C*at*^RN*Ai*^ and *W*^RN*Ai*^ dividing cells (mutant cells were identified by the absence of cytoplasmic GFP). The asterisk and horizontal bar indicate the time window for which the positions of the midbody in α-C*at*^RN*Ai*^ dividing cells significantly differ from the *W*^RN*Ai*^ dividing cells (*t*-test). (B-B’) Dlg::GFP (green in the top panel, white in the bottom panel) and MyoII::3mKate2 (red in the top panel) distributions in the dividing cell and its neighbours during cytokinesis in *Gli*^RN*Ai*^ clones (mutant cells were identified by the absence of cytoplasmic GFP). *Gli*^RN*Ai*^ cells are indicated by white dots. A full projection of the confocal z-section is shown. Apical-basal section of the cell in B at t=264 min (B’). White arrowheads, midbody; green arrowheads, the Dlg::GFP positive strands. (C) Graph of the midbody apical-basal positions (Mean±SEM) in *W*^RN*Ai*^ dividing cell, in wt dividing cell with one *Gli*^RN*Ai*^ and one wt neighbouring cell, in *Gli*^RN*Ai*^ dividing cell with one wt and one *Gli*^RN*Ai*^ neighbouring cells, and in *Gli*^RN*Ai*^ dividing cell surrounded by two *Gli*^RN*Ai*^ neighbouring cells (mutant cells were identified by the absence of cytoplasmic GFP). The asterisks and horizontal bars indicate the time window for which the positions of the midbody in mutant cells significantly differ from the control ones (*t*-test). (D,D’) Dlg::GFP (green in the top panel, white in the bottom panel) and MyoII::3mKate2 (red in the top panel) distributions in the dividing cell and its neighbours during cytokinesis in *aka*^RN*Ai*^ clones. *aka*^RN*Ai*^ cells are indicated by white dots. A full projection of the confocal z-section is shown. Apical-basal section of the cell in D at t=256 min (D’). White arrowheads: midbody. Green arrowheads indicate the Dlg::GFP positive strands. (E) Graph of the midbody apical-basal positions (Mean±SEM) in *W*^RN*Ai*^ dividing cell, in wt dividing cell with one *aka*^RN*Ai*^ and one wt neighbouring cells; in *aka*^RN*Ai*^ dividing cell with one wt and one *aka*^RN*Ai*^ neighbouring cells; and in *aka*^RN*Ai*^ dividing cell surrounded by two *aka*^RN*Ai*^ neighbouring cells (mutant cells were identified by the absence of cytoplasmic GFP). The asterisks and horizontal bars indicate the time window for which the positions of the midbody in mutant cells significantly differ from the controlled ones (*t*-test) Scale bars: 5 μm. *n*, cell numbers (A, C, E)

## Discussion

The coordination between *de novo* junction formation and cell division is a general feature of monolayered tissues (Denes et al., 2015; Firmino et al., 2016; Founounou et al., 2013; Herszterg et al., 2013; Higashi et al., 2016; Jinguji and Ishikawa, 1992; Lau et al., 2015; Morais-De-Sá and Sunkel, 2013; Pinheiro et al., 2017; Reinsch and Karsenti, 1994). The coordination of TCJ formation and cytokinesis leading to the formation of a neighbour-neighbour interface has been elegantly studied in the epithelium of the gastrulating *Xenopus* embryo. In this epithelium, a new junction is initially established between the neighbouring cells upon cytokinesis and 10 min after midbody formation the TCJ components Angulin-1 and Tricellulin are sequentially localized at the two cell vertices and prospective TCJs forming between the daughter and its neighbouring cells (Higashi et al., 2016). Here, we have explored the mechanisms of TCJ formation when a daughter-daughter interface is formed upon cell division. We found that the *de novo* TCJ assembly occurs during a stereotypical midbody movement, which is associated with the disentanglement of the neighbouring cell membranes from the daughter cell membranes and which depends on the activity of TCJ proteins.

Midbody dynamics have been studied in the context of cell division orientation, cell fate specification, dorsal-ventral axis specification and ciliogenesis as well as tumorigenesis (Bernabé-Rubio et al., 2016; Dubreuil et al., 2007; Ettinger et al., 2011; Kuo et al., 2011; Singh and Pohl, 2014). The mechanisms of midbody displacements have in particular been explored during dorsal-ventral axis specification in *C. elegans* where its movement is regulated by cortical flows (Singh and Pohl, 2014). A general feature of epithelial cells is the apical positioning of the midbody following cytokinesis (Firmino et al., 2016; Founounou et al., 2013; Guillot and Lecuit, 2013; Herszterg et al., 2013; Higashi et al., 2016; Jinguji and Ishikawa, 1992; Morais-De-Sá and Sunkel, 2013; Pinheiro et al., 2017; Reinsch and Karsenti, 1994); a position which is shown to be critical for tissue organisation (Morais-De-Sá and Sunkel, 2013). By performing a prolonged time-lapse analysis of the midbody dynamics, our work reveals the existence of a stereotypic midbody apical to basal displacement, which correlates with AJ, SJ and TCJ formation and which mainly depends on the Gli and Aka TCJ proteins. At least, two non-mutually exclusive mechanisms can be proposed to explain the midbody basal displacement. The displacement might by powered by the incorporation of junctional or cortical proteins in the newly formed apical interfaces; the accumulation of which might push the midbody to a more basal position by, for example promoting the adhesion between the two daughter cells. Such a mechanism requires to locally target the delivery of cortical or transmembrane proteins apical to the midbody. This mechanism is also consistent with our observation that the initial basal movement is delayed in the absence of the α-Catenin function, which is necessary for apical AJ formation. The second mechanism might be more actively regulated by midbody components. Indeed, the midbody is enriched in actin and microtubule molecular motors that are inherited upon the closure of the cytokinetic ring (Glotzer, 2017). These motors might displace the midbody along the cortex of the daughter cell membranes.

The formation of a 4-cell structure between the daughter and neighbour cells is a general feature of epithelial cytokinesis that is most likely instrumental to maintain epithelial barrier function. Previous works combined with our current study establish that the resolution of the 4-cell structure depends on at least two distinct and sequential steps. First, Rac GTPase and Arp2/3 dependent F-actin polymerization localized around the midbody promotes the neighbouring cell membrane withdrawal at the level of the AJs (Herszterg et al., 2013 and unpublished). The second step depends on the Gli and Aka TCJ proteins and ensures that the daughter and neighbouring cells are disengaged along their basal-lateral membrane. While the full characterization of Aka and Gli functions during this process requires deciphering the mechanisms promoting midbody movement, our work uncovers an unexpected role for TCJ proteins in the regulation of cytokinesis.

Together, our work illustrates the existence of an intricate interplay between midbody dynamics and the *de novo* TCJ formation and it unveils an additional step during cytokinesis necessary for the disengagement of the neighbouring cells from the midbody in epithelial tissue. Hence, our work opens novel avenues for the exploration of the mechanisms that ensure the coordination of cytokinesis and epithelial polarity establishment during division; a coordination allowing the epithelium to fulfil its basic physiological function: forming a paracellular barrier while being remodelled by proliferation during development, tissue homeostasis or regeneration.

## Materials and Methods

### Fly stocks and genetics

*Drosophila melanogaster* stocks are listed in Supplementary Table 1. Flies were crossed and experiments were performed at 25°C. Loss of function experiments were carried out using the FLP/FRT techniques (Bosveld et al., 2016a; Pinheiro et al., 2017; Xu and Rubin, 1993). Somatic clones were induced in the second instar larval stage by heat-shock at 37°C. The analyses were carried out 2-4 days after clone induction in 15-25 hours after pupa formation (hAPF) old pupae.

### Live imaging microscopy and photoactivation

Pupae were prepared for live imaging as described previously (Bosveld et al., 2016b). All experiments were performed during the first round of cell divisions in the anterior-central region of the notum tissue. The basal movement of the midbody is similar in all cells within this region (not shown) and we focused on the dynamics of the midbody in epithelial cells, fated to become tendon epithelial cells because it greatly facilitated the observations of the SJs and TCJs in 2D projections. In the analyses (unless mentioned otherwise), the time, t=0, was set at midbody formation. Samples were imaged at 25°C with an inverted confocal spinning disk microscope (Carl Zeiss), using 63x NA1.4 OIL DICII PL APO or 100x NA1.4 OIL DIC N2 PL APO VC objectives and either a CMOS (Hamamtsu) camera or EM-CCD (Roper) Camera. Photoactivation was performed with a two-photon Ti:Sapphire laser (Mai Tai DeepSee, Spectra Physics) at 820 nm with < 100 fs pulses typically set at 7% power, mounted on a confocal microscope (LSM880, Carl Zeiss) with a 63x NA 1.4 OIL DICII PL APO objective.

### Image quantifications

Segmentation of the PH::GFP signal was achieved using Imaris by manually drawing the contour of each cell and the position of the midbody. Midbody apical-basal movement was quantified by tracking the MyoII::3mKate2 labelled midbody position using Fiji (Schindelin et al., 2012). The midbody positions were quantified relative to its initial position of formation. The midbody formation position is located on average 1.24±0.06 μm in *W*^RN*Ai*^ (*n*=38) and 1.50±0.07 μm in FRT40A control cells (*n*=28), below the plane of the epithelial MyoII circumference belt. The position of midbody formation is similar in most experimental conditions analysed with the exception of *a-Cat*^RN*Ai*^ (0.97±0.10 μm, *n*=15 *p*<0.046), *aliX^1^* (1.64±0.10 μm, *n*=22,*p*<0.002), *Gli^dv3^* (1.81 μm±0.07 μm, *n*=65,*p*<0.01) and *aka^1,200^* (2.31 μm±0.12, *n*=21,*p*<0.0001) dividing cells. The difference in *a-Cat*^RN*Ai*^, *Gli* and *aka* dividing cells could reflect defects in apical-basal polarity of the cells. The defects observed in *aliX^1^* cells are more difficult to interpret since the positions of the midobody were normal in *aliX^1^/aliX^3^* and in *aliX^3^* dividing cells.

All images, unless specified otherwise, represent maximal intensity projections of the z-stacks. Apical-basal (side views) section for which no dashed line is shown in the top view images correspond to images that were rotated in 3D using the TransformJ Fiji plugin to provide better enface views of the newly forming SJ and TCJs. Figures were created using Fiji and Abode Illustrator.

Detachement of the Dlg::GFP positive strands was analysed between 0 and 180 min post midbody formation in *W*^RN*Ai*^, *Gli*^RN*Ai*^, *aka*^RN*Ai*^, *Gli^dv3^*, *aka^1,200^* and control dividing cells remaining in contact with *W*^RN*Ai*^, *Gli*^RN*Ai*^, *aka*^RN*Ai*^, *Gli^dv3^*, *aka^1,200^* and control neighbouring cells, respectively, during the whole period. In some cells, the Dlg::GFP positive strands were located just below the junctions, precluding the characterization of their detachment from the midbody. We therefore excluded these cells from the analysis corresponding to 9 out of 30 *Gli*^RN*Ai*^ cells, 4 out of 24 *aka*^RN*Ai*^ cells, 8 out of 24 *W*^RN*Ai*^ cells, 40 out of 65 *Gli^dv3^*cells, 6 out of 21 *aka^1,200^* cells and 71 out of 89 FRT40A control cells.

### Statistics

Statistical analyses were performed using the Student *t*-test. Significant differences for midbody apical-basal positions are reported if the positions of 3 consecutive time points were significantly (*p*<0.05) different from the control cells. Error bars reported in the manuscript are standard error of the mean, SEM.

## Acknowledgments

We thank Vanessa Auld (University of British Columbia, Vancouver, Canada), Kaisa Haglund (Oslo University Hospital, Oslo, Norway), Jean-René Huynh (Collège de France, Paris, France), Stefan Luschnig (University of Münster, Münster, Germany), Benedicte Sansson (Cambridge University, Cambridge, UK), Andreas Wodarz (University of Göttingen, Göttingen, Germany), the Transgenic RNAi Project at Harvard Medical School and VDRC, the Kyoto and Bloomington Stock Centers for reagents; the Developmental Biology Curie imaging facility P1CT-IBiSA@DBB for help with microscopy; F. Agnes, F. Di Pietro, A. Maugarny-Cales and E. Morais-de-Sá for comments on the manuscript.

## Author contributions

Conceptualization: Z.W., F.B., Y.B.; Methodology: Z.W., F.B., Y.B.; Validation: Z.W., Y.B.; Formal analysis: Z.W., F.B., Y.B.; Investigation: Z.W.; Data curation: Z.W.; Writing – original draft: Y.B.; Writing - review & editing: Z.W., F.B., Y.B.; Visualization: Z.W., F.B., Y.B.; Supervision: F.B., Y.B.; Project administration: Y.B.; Funding acquisition: Y.B.

---

**Supplementary Table 1:** Alleles and transgenes used in this study.

| ***Drosophila* stock** | **Reference or Source** |
| --- | --- |
| E-Cad::3mKate2 | Pinheiro et al., 2017 |
| MyoII::3mKate2 (Sqh) | Pinheiro et al., 2017 |
| UAS-Lifeact::GFP | Bloomington Stock Center |
| Lac::YFP | Lye et al., 2014 |
| Dlg::GFP | Morin et al., 2001 |
| Dlg::TagRFP | Pinheiro et al., 2017 |
| UAS-shrb::GFP | Matias et al., 2015 |
| Gli::YFP | Lye et al., 2014 |
| Aka::GFP | Byri et al., 2015 |
| Jup::GFP | Morin et al., 2001 |
| Nrg::GFP | Morin et al., 2001 |
| α-ATPase::GFP | Morin et al., 2001 |
| UAS-Tub::PAFP | Bloomington Stock Center |
| UAS-PH::GFP | von Stein et al., 2005 |
| Actin5C-FRT-y<sup>+</sup>-FRT-Gal4 | Bloomington Stock Center |
| Tub-FRT-GFP-FRT-Gal4 | Bloomington Stock Center |
| FRT42D nls::GFP | Bloomington Stock Center |
| Actin5C-Gal4 | Bloomington Stock Center |
| alix<sup>1</sup> FRT82B | Eikenes et al., 2015 |
| alix<sup>3</sup> FRT82B | Eikenes et al., 2015 |
| aka<sup>L200</sup> FRT40A | Byri et al., 2015 |
| Gli<sup>dv3</sup> FRT40A | Venema et al., 2004 |
| FRT40A | Bloomington Stock Center |
| UAS-shrb<sup>TRiP.HMS01767</sup> | Bloomington Stock Center |
| UAS-Gli<sup>TRiP.HM05262</sup> | Bloomington Stock Center |
| UAS-aka<sup>GD52608</sup> | VDRC |
| UAS-α-Cat<sup>KK107916</sup> | VDRC |
| UAS-w<sup>TRiP.HMS00045</sup> | Bloomington Stock Center |

## Supplementary Movies

**Supplementary Movie 1. Upon cytokinesis AJ formation precedes SJ formation and a 4-cell structure is created.**

Confocal time-lapse movie of Dlg::GFP (green) and E-Cad::3mKate2 (red) distributions in the dividing cell and its neighbours during cytokinesis. E-Cad::3mKate2 signal was projected at the level of the AJ and Dlg::GFP was projected at the level of the SJ to highlight their respective dynamics. Green arrowheads, deforming septate junction marked by Dlg::GFP at the 4-cell structure formed at the interface between the two daughter cells and neighbouring cells; red arrowheads, forming AJ between the two daughter cells.

Scale bar: 5 μm.

**Supplementary Movie 2. Dynamics of Dlg::GFP during the formation of the Dlg::GFP positive strands.**

Confocal time-lapse movie of Dlg::GFP (green) and MyoII::3mKate2 (red) distributions in the dividing cell (asterisk) and its neighbours during cytokinesis. A full projection of the confocal z-section is shown. White arrowheads, midbody; green arrowheads, presumptive TCJ strands forming between the apical SJ and the midbody; open green arrowheads, TCJ strands unattached to the midbody.

Scale bar: 5 μm.

**Supplementary Movie 3. Gli positive TCJ strands form and are attached to the midbody as it descends.**

Confocal time-lapse movie of Gli::YFP (green) and MyoII::3mKate2 (red) distributions in a dividing cell (asterisk). A full projection of the confocal z-section is shown. White arrowheads, midbody; green arrowheads Gli::YFP strands extending towards and apposed to the midbody; open green arrowheads, Gli::YFP positive strands detached from the midbody.

**Supplementary Movie 4. The presumptive TCJ stands remain attached to the midbody upon Gli knock-down.**

Confocal time-lapse movie of Dlg::GFP (green) and MyoII::3mKate2 (red) distributions in the dividing cell (asterisk) and its neighbours during cytokinesis in *Gli*^RN*Ai*^ flip-out clones. *Gli*^RN*Ai*^ dividing cell (asterisk) and the 2 neighbour cells marked by white dots (mutant cells were identified by the absence of cytoplasmic GFP). A full projection of the confocal z-section is shown. White arrowhead: midbody. Green arrowheads indicate the Dlg::GFP strands that failed to detach from the midbody.

## References

Bernabé-Rubio, M., Andrés, G., Casares-Arias, J., Fernández-Barrera, J., Rangel, L., Reglero-Real, N., Gershlick, D. C., Fernández, J. J., Millán, J., Correas, I., et al. (2016). Novel role for the midbody in primary ciliogenesis by polarized epithelial cells.. J. Cell Biol. 214, 259–273.

Bosveld, F., Markova, O., Guirao, B., Martin, C., Wang, Z., Pierre, A., Balakireva, M., Gaugue, I., Ainslie, A., Christophorou, N., et al. (2016a). Epithelial tricellular junctions act as interphase cell shape sensors to orient mitosis. Nature 530, 495–498.

Bosveld, F., Guirao, B., Wang, Z., Rivière, M., Bonnet, I., Graner, F. and Bellaïche, Y. (2016b). Modulation of junction tension by tumor suppressors and proto-oncogenes regulates cell-cell contacts. Development 143, 623–634.

Byri, S., Misra, T., Syed, Z. A., Bätz, T., Shah, J., Boril, L., Glashauser, J., Aegerter-Wilmsen, T., Matzat, T., Moussian, B., et al. (2015). The Triple-Repeat Protein Anakonda Controls Epithelial Tricellular Junction Formation in Drosophila. Dev. Cell 33, 535–548.

Capalbo, L., Montembault, E., Takeda, T., Bassi, Z. I., Glover, D. M. and D’Avino, P. P. (2012). The chromosomal passenger complex controls the function of endosomal sorting complex required for transport-III Snf7 proteins during cytokinesis. Open Biol. 2, 120070.

Denes, A. S., Kanca, O., Affolter, M., Vincent, S., Affolter, M. and Schuh, R. (2015). A cellular process that includes asymmetric cytokinesis remodels the dorsal tracheal branches in Drosophila larvae. Development 142, 1794–805.

Dubreuil, V., Marzesco, A. M., Corbeil, D., Huttner, W. B. and Wilsch-Bräuninger, M. (2007). Midbody and primary cilium of neural progenitors release extracellular membrane particles enriched in the stem cell marker prominin-1. J. Cell Biol. 176, 483–495.

Eikenes, Å. H., Malerød, L., Christensen, A. L., Steen, C. B., Mathieu, J., Nezis, I. P., Liestøl, K., Huynh, J. R., Stenmark, H. and Haglund, K. (2015). ALIX and ESCRT-III Coordinately Control Cytokinetic Abscission during Germline Stem Cell Division In Vivo. PLoS Genet. 11, e1004904.

Ettinger, A. W., Wilsch-Bräuninger, M., Marzesco, A. M., Bickle, M., Lohmann, A., Maliga, Z., Karbanová, J., Corbeil, D., Hyman, A. A. and Huttner, W. B. (2011). Proliferating versus differentiating stem and cancer cells exhibit distinct midbody-release behaviour. Nat. Commun. 2, 503.

Firmino, J., Rocancourt, D., Saadaoui, M., Moreau, C. and Gros, J. (2016). Cell Division Drives Epithelial Cell Rearrangements during Gastrulation in Chick. Dev. Cell 36, 249–261.

Founounou, N., Loyer, N. and Le Borgne, R. (2013). Septins Regulate the Contractility of the Actomyosin Ring to Enable Adherens Junction Remodeling during Cytokinesis of Epithelial Cells. Dev. Cell 24, 242–255.

Glotzer, M. (2017). Cytokinesis in metazoa and fungi. Cold Spring Harb. Perspect. Biol. 9, a022343.

Guillot, C. and Lecuit, T. (2013). Adhesion Disengagement Uncouples Intrinsic and Extrinsic Forces to Drive Cytokinesis in Epithelial Tissues. Dev. Cell 24, 227–241.

Herszterg, S., Leibfried, A., Bosveld, F., Martin, C. and Bellaiche, Y. (2013). Interplay between the Dividing Cell and Its Neighbors Regulates Adherens Junction Formation during Cytokinesis in Epithelial Tissue. Dev. Cell 24, 256–270.

Herszterg, S., Pinheiro, D. and Bellaïche, Y. (2014). A multicellular view of cytokinesis in epithelial tissue. Trends Cell Biol. 24, 285–293.

Higashi, T. and Miller, A. L. (2017). Tricellular junctions: how to build junctions at the TRICkiest points of epithelial cells. Mol. Biol. Cell 28, 2023–2034.

Higashi, T., Arnold, T. R., Stephenson, R. E., Dinshaw, K. M. and Miller, A. L. (2016). Maintenance of the Epithelial Barrier and Remodeling of Cell-Cell Junctions during Cytokinesis. Curr. Biol. 26, 1829–1842.

Jinguji, Y. and Ishikawa, H. (1992). Electron microscopic observations on the maintenance of the tight junction during cell division in the epithelium of the mouse small intestine. Cell Struct. Funct. 17, 27–37.

Kuo, T. C., Chen, C. T., Baron, D., Onder, T. T., Loewer, S., Almeida, S., Weismann, C. M., Xu, P., Houghton, J. M., Gao, F. B., et al. (2011). Midbody accumulation through evasion of autophagy contributes to cellular reprogramming and tumorigenicity. Nat. Cell Biol. 13, 1214–1223.

Lau, K., Tao, H., Liu, H., Wen, J., Sturgeon, K., Sorfazlian, N., Lazic, S., Burrows, J. T. A., Wong, M. D., Li, D., et al. (2015). Anisotropic stress orients remodelling of mammalian limb bud ectoderm. Nat. Cell Biol. 17, 569–579.

Lye, C. M., Naylor, H. W. and Sanson, B. (2014). Subcellular localisations of the CPTI collection of YFP-tagged proteins in Drosophila embryos. Development 141, 4006–4017.

Matias, N. R. eis, Mathieu, J. and Huynh, J. R. (2015). Abscission is regulated by the ESCRT-III protein shrub in Drosophila germline stem cells. PLoS Genet. 11, e1004653.

Mierzwa, B. and Gerlich, D. W. (2014). Cytokinetic Abscission: Molecular Mechanisms and Temporal Control. Dev. Cell 31, 525–538.

Morais-De-Sá, E. and Sunkel, C. (2013). Adherens junctions determine the apical position of the midbody during follicular epithelial cell division. EMBO Rep. 14, 696–703.

Morin, X., Daneman, R., Zavortink, M. and Chia, W. (2001). A protein trap strategy to detect GFP-tagged proteins expressed from their endogenous loci in Drosophila. Proc. Natl. Acad. Sci. 98, 15050–15055.

Pinheiro, D., Hannezo, E., Herszterg, S., Bosveld, F., Gaugue, I., Balakireva, M., Wang, Z., Cristo, I., Rigaud, S. U., Markova, O., et al. (2017). Transmission of cytokinesis forces via E-cadherin dilution and actomyosin flows. Nature 545, 103–107.

Reinsch, S. and Karsenti, E. (1994). Orientation of spindle axis and distribution of plasma membrane proteins during cell division in polarized MDCKII cells. J. Cell Biol. 126, 1509–1526.

Schindelin, J., Arganda-Carreras, I., Frise, E., Kaynig, V., Longair, M., Pietzsch, T., Preibisch, S., Rueden, C., Saalfeld, S., Schmid, B., et al. (2012). Fiji: An open-source platform for biological-image analysis. Nat. Methods 9, 676–682.

Singh, D. and Pohl, C. (2014). Coupling of Rotational Cortical Flow, Asymmetric Midbody Positioning, and Spindle Rotation Mediates Dorsoventral Axis Formation in C. elegans. Dev. Cell 28, 253–267.

Venema, D. R., Zeev-Ben-Mordehai, T. and Auld, V. J. (2004). Transient apical polarization of Gliotactin and Coracle is required for parallel alignment of wing hairs in Drosophila. Dev. Biol. 275, 301–314.

von Stein, W., Ramrath, A., Grimm, A., Müller-Borg, M. and Wodarz, A. (2005). Direct association of Bazooka/PAR-3 with the lipid phosphatase PTEN reveals a link between the PAR/aPKC complex and phosphoinositide signaling. Development 132, 1675–1686.

Xu, T. and Rubin, G. M. (1993). Analysis of genetic mosaics in developing and adult Drosophila tissues. Development 117, 1223–1237.

